# Circulating Microbial DNA as a Potential Cancer Biomarker: Technical Challenges and Controlled Evaluation

**DOI:** 10.64898/2026.08.17.745053

**Authors:** Daisy Fry Brumit, Daniel Bsteh, Shan Sun, Jesse A. Goodrich, Tanya L. Alderete, Michael A. Liss, Amir Goldkorn, Anthony A. Fodor

**Affiliations:** Department of Bioinformatics and Genomics, University of North Carolina at Charlotte, Charlotte, North Carolina; Department of Environmental Health and Engineering, Bloomberg School of Public Health, Johns Hopkins University, Baltimore, Maryland; Division of Medical Oncology, Department of Medicine, Keck School of Medicine of University of Southern California, Los Angeles, California; USC Norris Comprehensive Cancer Center, Los Angeles, California; Department of Population and Public Health Sciences, Keck School of Medicine, University of Southern California, Los Angeles, California; Department of Urology, Center for Microbiome Innovation, University of California San Diego, San Diego, California

**Keywords:** liquid biopsy, cancer microbiome, prostate cancer, genitourinary malignancy, plasma cfDNA, circulating microbial DNA, metagenomic sequencing

## Abstract

**Background:** Circulating microbial DNA (cmDNA) has been proposed as a non-invasive cancer biomarker, but most evidence comes from cancer-sequencing datasets not designed for microbial analysis and lacking contamination controls. Whether reported signatures reflect biology or artifact is unclear in low-biomass specimens, where standard taxonomic pipelines are prone to systematic error.

**Methods:** In a tightly controlled pilot study of metastatic castration-resistant prostate cancer, we profiled plasma cell-free DNA (cfDNA) and buffy-coat genomic DNA (gDNA) from two patients and two healthy volunteers alongside mock blood-draw and reagent controls, each with and without host-DNA depletion. Reads were classified with Kraken2/Bracken and, independently, with the marker-gene classifier MetaPhlAn. As informatics controls, reads were per-base shuffled to randomize nucleotide order while preserving read length and guanine-cytosine (GC) content, and purely synthetic reads were generated from a four-base process matched only to an aggregate GC target; both were classified identically. Genus abundances were regressed against Kraken2 database k-mer representation and against GC content.

**Results:** Across 40 samples, Kraken2 reported several thousand genera, samples clustered by specimen type in principal-coordinate analysis (PCoA), and pooled genus counts correlated strongly with a published cancer-microbiome catalog (The Cancer Genome Atlas lung adenocarcinoma, TCGA-LUAD; Spearman ρ = 0.81 over 282 shared genera), a pattern readily interpreted as biological signal. However, these observations were also made in per-base shuffling, which preserves GC content and length but destroys all biological sequence: shuffled reads were still abundantly classified, still clustered by specimen type, and still correlated with the catalog (ρ ≈ 0.7), as did every sample group, including pure reagent controls. Genus counts scaled tightly with each genus’s k-mer representation in the Kraken2 database on real (r² = 0.74) and shuffled (r² = 0.85) reads, and the same dependence appeared in the independent published cohort. Purely synthetic reads carrying no information beyond an aggregate GC target reproduced much of the cross-cohort agreement (synthetic TCGA-LUAD ρ = 0.61 versus 0.81 for real reads; significant in 27 of 33 TCGA cancers), and replicate shuffles of a low-GC versus a high-GC plasma sample, for which the true difference is zero, produced spurious significant differences in about 46% of genera. Regressing observed counts against the shuffled baseline left 23 genera above the artifact floor at 5% false discovery rate (FDR), nearly all known kit contaminants, control-enriched viruses, or very-low-abundance taxa; a four-criterion validity filter reduced thousands of Kraken2 genera to a single defensible candidate, *Klebsiella*.

**Conclusions:** Much of the apparent cmDNA structure, including its agreement with a published cancer-microbiome catalog, is explained by base composition and reference-database architecture rather than authentic biology, and short-read k-mer pipelines cannot separate the two on their own. We find little positive evidence of an authentic circulating microbial signal, though our small sample cannot prove its absence. To limit false discovery in low-biomass metagenomics, we recommend specimen-matched negative controls, corroboration with a conservative second classifier, per-base shuffling (with GC-matched synthetic reads as a stricter floor), and GC-aware analysis.

**Graphical Abstract:** Pipeline pitfalls in blood microbial-DNA assays. cmDNA from low-biomass blood is vulnerable to three errors across the workflow (collection, processing, library preparation and sequencing, informatics): environmental (non-blood) contamination, kit and reagent contamination, and taxonomic misclassification. The corresponding safeguards are mock and specimen-matched negative controls, a literature sweep for known contaminants, and corroboration of k-mer output with a marker-gene classifier and a shuffle-based artifact control.

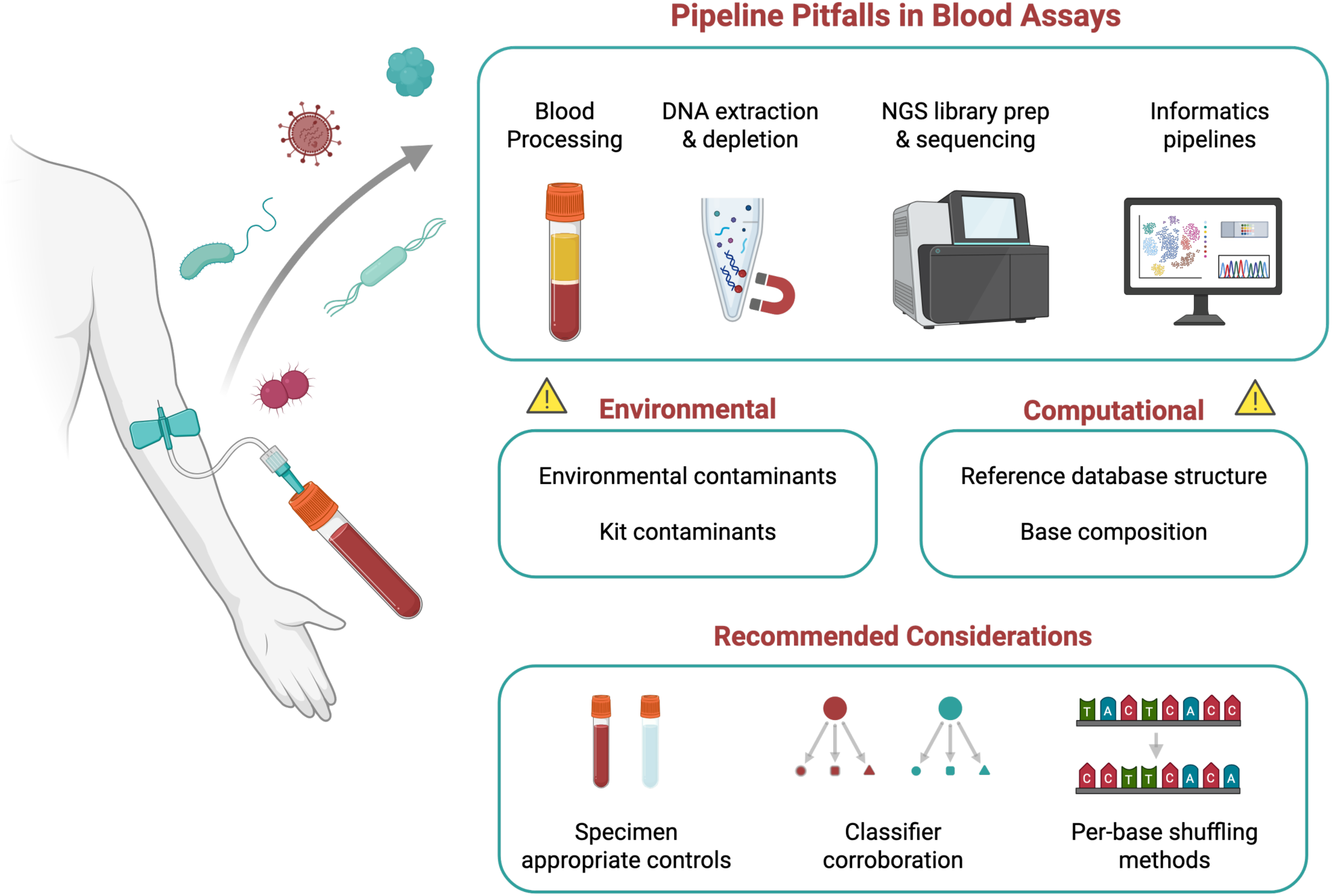

## Introduction

The human microbiome is increasingly recognized as a modulator of cancer biology, with dysbiosis linked to tumorigenesis, immune evasion, and therapy response. This interest has extended beyond gastrointestinal malignancies to genitourinary (GU) tumors, where tissue-associated and liquid-biopsy microbial signatures are being explored. Systematic reviews report altered urinary microbial compositions in bladder, prostate, and renal cell carcinoma, with associations to stage, recurrence, and grade [1,2]. In prostate cancer, a meta-analysis of 2,195 patients found certain GU taxa depleted relative to controls, although urine-only analyses did not reach significance, underscoring methodological heterogeneity [2]. Beyond the gut, intratumoral, and urinary microbiomes, circulating microbial DNA (cmDNA) offers a further non-invasive source of microbial signal. This cmDNA can be found in blood plasma and other fluids and blood microbial signatures have been reported in conditions such as smoking and chronic kidney disease [7,8].

Blood is a non-invasive, repeatable route to biomarker discovery, enabling serial sampling across disease progression and treatment. Yet basic questions remain open, such as whether plasma is the optimal analyte or whether peripheral blood mononuclear cells (PBMCs) or extracellular vesicles yield stronger or more stable readouts. Another unresolved question is whether physiological variables which influence circulating nucleic acid dynamics, such as circadian rhythm and feeding state, matter here. Compounding this, microbial-detection algorithms developed for the gut microbiome or other microbially rich specimens can exhibit substantial systematic biases on low-biomass fluids such as plasma, generating artifactual signatures [3].

Most cmDNA studies to date rely on repurposed cancer-sequencing datasets not designed for microbial analysis and lacking contamination controls. A reported pan-cancer blood and tissue microbiome from The Cancer Genome Atlas (TCGA), and diagnostic classifiers built from it, drew wide interest [9], but independent reanalyses have questioned how much reflects biology rather than contamination and bioinformatic artifact [10,11]. These differences in specimen handling, sequencing, and analysis have fueled debate over the validity of reported microbial cfDNA signatures in cancer.

To address this, we designed a controlled pilot study to identify which blood analyte gives the most reliable microbial signal and to characterize contamination introduced during collection, processing, and sequencing. As shown below, the compositional structure a standard short-read pipeline recovers from blood is driven largely by GC content and reference-database architecture rather than microbial biology, and a sequence-shuffling control exposes this artifact. This small but highly controlled study confronts the challenges of low-biomass profiling, provides a rigorously controlled dataset for evaluating current tools, and lays a foundation for larger efforts to define authentic circulating microbial signatures. We further show that GC differences between samples inflate false-positive rates in per-taxon testing, which argues for judging required effect sizes against the shuffle-derived artifact floor.

## Methods

### Study design and specimens

The study evaluated the optimal blood specimen type for microbial-signature detection in metastatic castration-resistant prostate cancer (mCRPC) while quantifying contamination (Figure 1). Blood was collected from two healthy volunteers (each in duplicate cell-free DNA Streck BCT tubes) and two patients with mCRPC treated at USC Norris Comprehensive Cancer Center; each volunteer was sampled in the morning and afternoon, so that time of day, health status, host depletion, and control type could be evaluated as factors of interest. To capture contamination from phlebotomy, skin, or collection materials, four mock blood draws were performed by perforating the skin with standard sterile technique but aspirating autoclaved water instead of blood. A commercial cfDNA reference material (SeraSeq Cell-Free Genomic Reference DNA) was spiked into the mock draws to provide input DNA, and the reference material alone was also sequenced to identify contaminants intrinsic to the control. Analyte factors were examined in parallel: plasma versus buffy coat, each with or without host-DNA depletion, for 40 samples in total. All participants provided written informed consent, and blood collection was performed under University of Southern California (USC) Institutional Review Board (IRB)-approved protocols HS-17-00639 (volunteers) and HS-11-00054 (patients).

**Figure 1.**
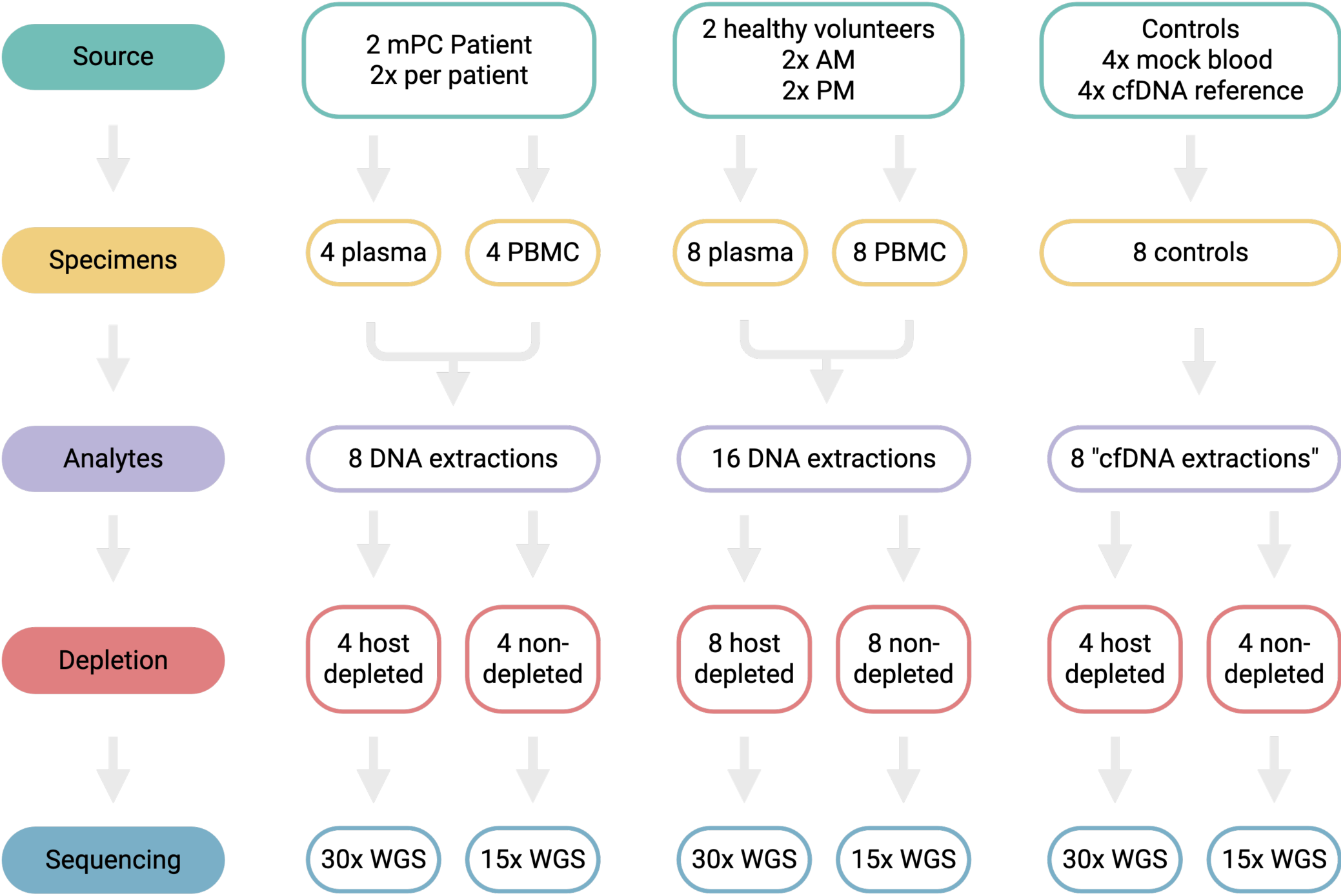
Study design. Blood from two metastatic castration-resistant prostate cancer (mPC) patients and two healthy volunteers (sampled morning, AM, and afternoon, PM), with contamination controls (four mock blood draws, four cell-free DNA reference-material controls). Specimens were split into plasma and buffy-coat (peripheral blood mononuclear cell, PBMC) fractions, extracted, prepared as host-depleted and non-depleted libraries, and sequenced by whole-genome sequencing (hWGS) at approximately 30× (non-depleted) or 15× (depleted), for 40 libraries. The design lets time of day, health status, host depletion, and control type be evaluated as factors of interest.

### Sample processing

Blood underwent double centrifugation to separate plasma and buffy coat. Plasma and mock controls were extracted for cfDNA with the nRichDX Revolution cfDNA Max Kit; buffy-coat genomic DNA (gDNA) was isolated with the NEB Monarch Genomic DNA Purification Kit.

### Library preparation and sequencing

Two replicates were prepared per sample type: one by direct library preparation, one after host (human) DNA depletion. Direct libraries used the NEBNext Ultra II DNA Library Prep Kit (NEB Cat. # E7645) with NEBNext Unique Dual Indexed unique molecular identifier (UMI) adapters (NEB Cat. # E7395); Human DNA-depleted libraries used the DNA Enrichment Kit (NEB Cat. # E2612), which removes CpG-methylated human DNA to enrich unmethylated microbial DNA. Libraries passed Qubit and Agilent BioAnalyzer quality control and were sequenced on an Illumina NovaSeq X Plus (PE150) to approximately 30× (non-depleted) or 15× (depleted) coverage.

### Metagenomic processing

Reads were trimmed with Trimmomatic 0.39 [17] (leading/trailing Q20, 4:20 sliding window, minimum length 50), and human reads removed with Hostile [18] (paired-end). Host-removed reads were profiled with two pipelines: Kraken2 [14] for taxonomic classification with genus-level abundance by Bracken [15], and, independently, MetaPhlAn [16] as a marker-gene estimate.

### Per-base shuffling control

To assess Kraken2-suite bias, host-removed reads were shuffled by randomizing base order within each read, preserving exact GC content and length while destroying biological sequence [3], and reclassified identically with Kraken2 and Bracken.

### Downstream analyses

Unshuffled and shuffled profiles were compared across specimen types. For each dataset, the most abundant taxa were retained and the rest pooled as “Other” in stacked bar plots ordered by specimen type, and the number of taxa above 0.01% mean relative abundance was recorded. Relative-abundance tables were visualized by PCoA on Bray-Curtis dissimilarities (R vegan). Per-taxon mean abundances were compared between predefined group pairs spanning clinical status, specimen type, host-depletion status, time of collection, and shuffling status, excluding taxa present in fewer than 1% of samples. To test whether detections tracked database representation, the k-mers each genus contributes to the Kraken2 standard database were counted, matched to mean genus counts, log₁₀-transformed, and regressed (R², slope reported

Linear models also tested PCoA axis loadings against GC content, and taxon abundance against database representation, for both datasets. Analyses used R (ape, vegan, lme4, lmerTest, RColorBrewer, ggrepel, tidyverse).

### Synthetic-read generation

As a stricter control, 10⁷ synthetic paired-end reads were generated by an independent Java program (L64X128MixRandom; seeds R1 = 20260618, R2 = 20260619). Each base was drawn independently with p(A) = p(T) = 0.271 and p(G) = p(C) = 0.229 (GC target 0.458); no Ns. Read lengths followed L = min(150, max(50, round(150 − E))), E ∼ Exponential(scale = 10.7), matching the post-trimming distribution (50 to 150 bp; observed mean 140 bp, synthetic mean 139.3 bp). R1 and R2 lengths were independent, with no insert or overlap structure; quality was Phred 40. Determinism was verified by sha256-matched regeneration of a 100,000-pair pilot.

### Synthetic-read classification

Synthetic reads were classified with the same Kraken2 standard database (build 2025-10-15; k = 35, minimizer 31) and software as the other libraries: Kraken2 2.1.3 (--paired --report-zero-counts --gzip-compressed --use-names; confidence 0.0, minimum hit groups 2) and Bracken 2.9 (-l G -r 100). All 17 database files were md5-identical to the study build, including hash.k2d (md5 e8336b184b83083e1b7ec149a0f047c2).

### Publicly available datasets

Study profiles were compared with the publicly available pan-cancer microbiome catalog of Sepich-Poore et al. [11], which reprocessed sequencing data from The Cancer Genome Atlas (TCGA) to derive decontaminated microbial read counts and extends the original TCGA blood-and-tissue microbiome analysis of Poore et al. [9]. We used their genus-level count table (Table S8; microbial counts derived by KrakenUniq classification of T2T-aligned TCGA whole-genome and whole-transcriptome sequencing, which spans 7,827 samples across the 33 TCGA cancer investigations listed in their Table S9. Every comparison was restricted to the 282 genera shared between that panel and our Kraken2/Bracken output, and per-genus counts were pooled within each cohort, so the resulting cohort-level correlations are upper bounds that suppress within-sample variance. TCGA lung adenocarcinoma (TCGA-LUAD; n = 395 samples) is the running example throughout; the choice is arbitrary, and the same agreement holds across the large majority of the 33 cohorts (Figure 7). Two further public resources were used as distributed: the Kraken2 standard reference database (build 2025-10-15), which both classified all libraries and provided the per-genus k-mer inventory, and the reference index bundled with Hostile for human-read removal. The TCGA counts derive largely from tumor and adjacent-tissue sequencing rather than low-biomass plasma, so the cross-cohort comparison probes shared classifier behavior rather than like-for-like biology.

### Data and code availability

Sequencing data is accessible as a BioProject with accession number PRJNA1466274. Code associated with this project can be found on GitHub at https://github.com/afodor/daisy-cmdna-verification.

## Results

### Standard analysis shows apparent clustering by specimen type and agreement with a published cancer microbiome

Kraken2 initially identified approximately 50 genera above 0.01% mean relative abundance across all samples. Buffy coats were dominated by *Bradyrhizobium*, controls carried inflated *Lymphocryptovirus*, and remaining samples retained a high proportion of reads classified as *Homo* even after host-read removal, though Figure 2A shows only microbial genera and not the residual human fraction. PCoA supported clear clustering by specimen type and separation from controls (Figure 2B), and over the 282 genera shared with a published lung adenocarcinoma catalog [11], our plasma samples correlated strongly (Figure 2C). Taken at face value, this could be read as distinct microbial communities, with patient samples reproducing an existing cancer count table, and would seem to confirm a robust, broadly distributed blood microbiome reproducible across cohorts and pipelines. The analyses below test whether these results instead reflect the classifier and its reference database.

**Figure 2.**
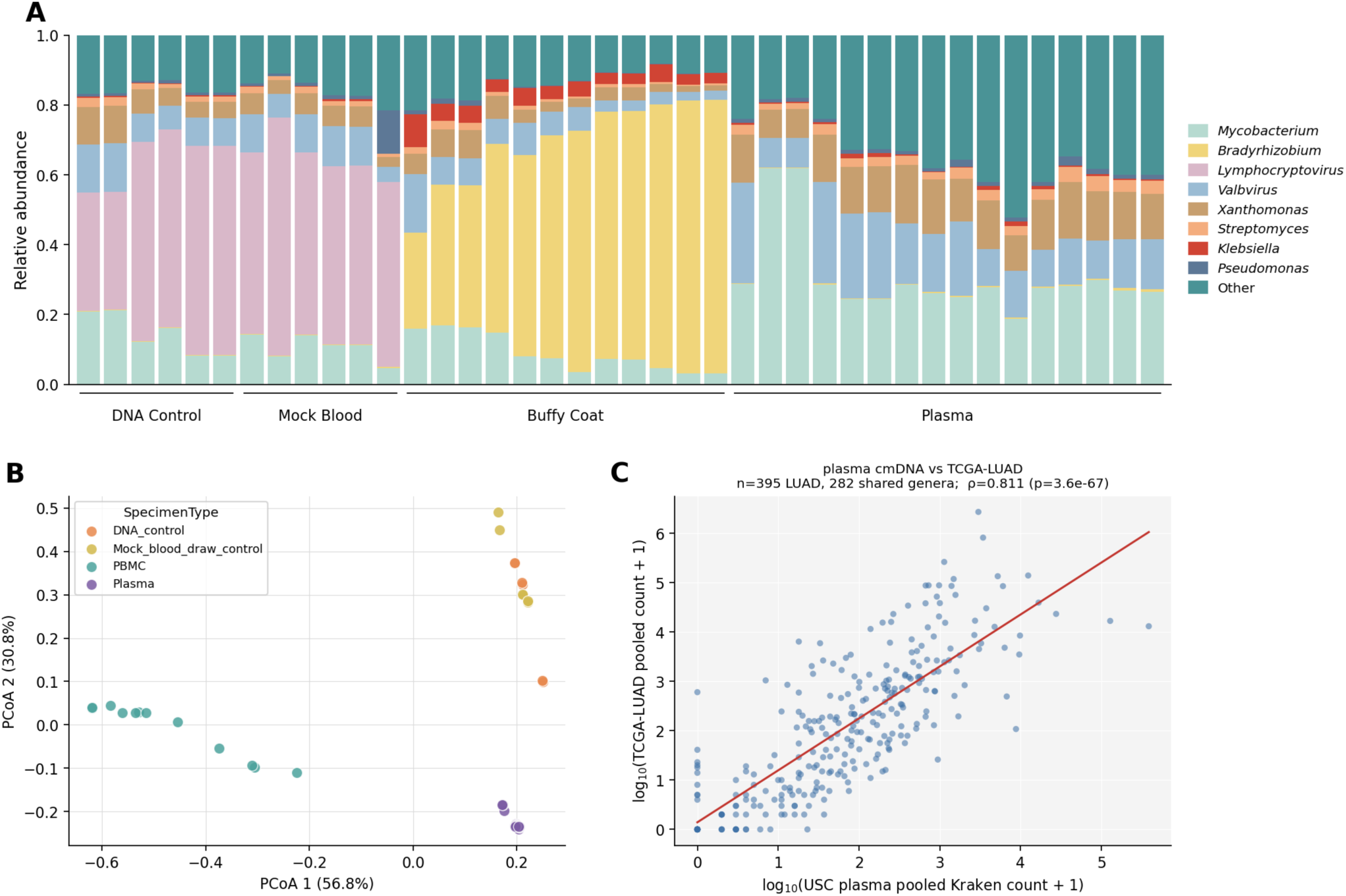
Genus-level results on unshuffled reads (40 libraries). (A) Relative-abundance stacked bars grouped by specimen type; the eight most abundant genera are colored, the rest pooled as “Other.” Buffy coats are dominated by *Bradyrhizobium*; other libraries are driven by *Valbvirus* and *Mycobacterium*, with *Lymphocryptovirus* in DNA-control and mock-blood samples and *Xanthomonas* in plasma. (B) PCoA on Bray-Curtis distance separates samples almost entirely by specimen type along PCoA 1 (56.8%), with DNA controls, mock draws, buffy coats, and plasma in largely non-overlapping regions. (C) Pooled per-genus Kraken2 counts for the 16 plasma libraries versus pooled TCGA-LUAD counts (n = 395; Sepich-Poore et al., Table S8) over 282 shared genera; Spearman ρ = 0.81, p = 3.6 × 10⁻⁶⁷; red line, ordinary least-squares (OLS) fit in log₁₀ space. [Figure to be regenerated to remove the “Daisy” axis label in panel C.]

### Shuffling sequences also yields separation by specimen type, driven by reference-database structure

Because compositions clustered by specimen type and correlated with an unrelated dataset (our plasma versus TCGA-LUAD), we probed their robustness. As a control, each host-removed read was shuffled to randomize nucleotide order while preserving GC content and length [3]. Shuffled sequences were still abundantly classified by Kraken2 (Figure 3A), still separated by specimen type (Figure 3B), and still correlated with the reported lung-cancer microbiome (Figure 3C), though less strongly than the unshuffled sequences (Figure 2C). Repeating the comparison against each published cancer type individually, the shuffled classifications correlated with every cancer, with per-cancer R² reaching 0.4 to 0.5 (Figure 4). Thus, the per-genus quantities Kraken2 reports, in our hands and in published low-biomass surveys, are partly a property of how the reference database is populated rather than of the microbial community.

**Figure 3.**
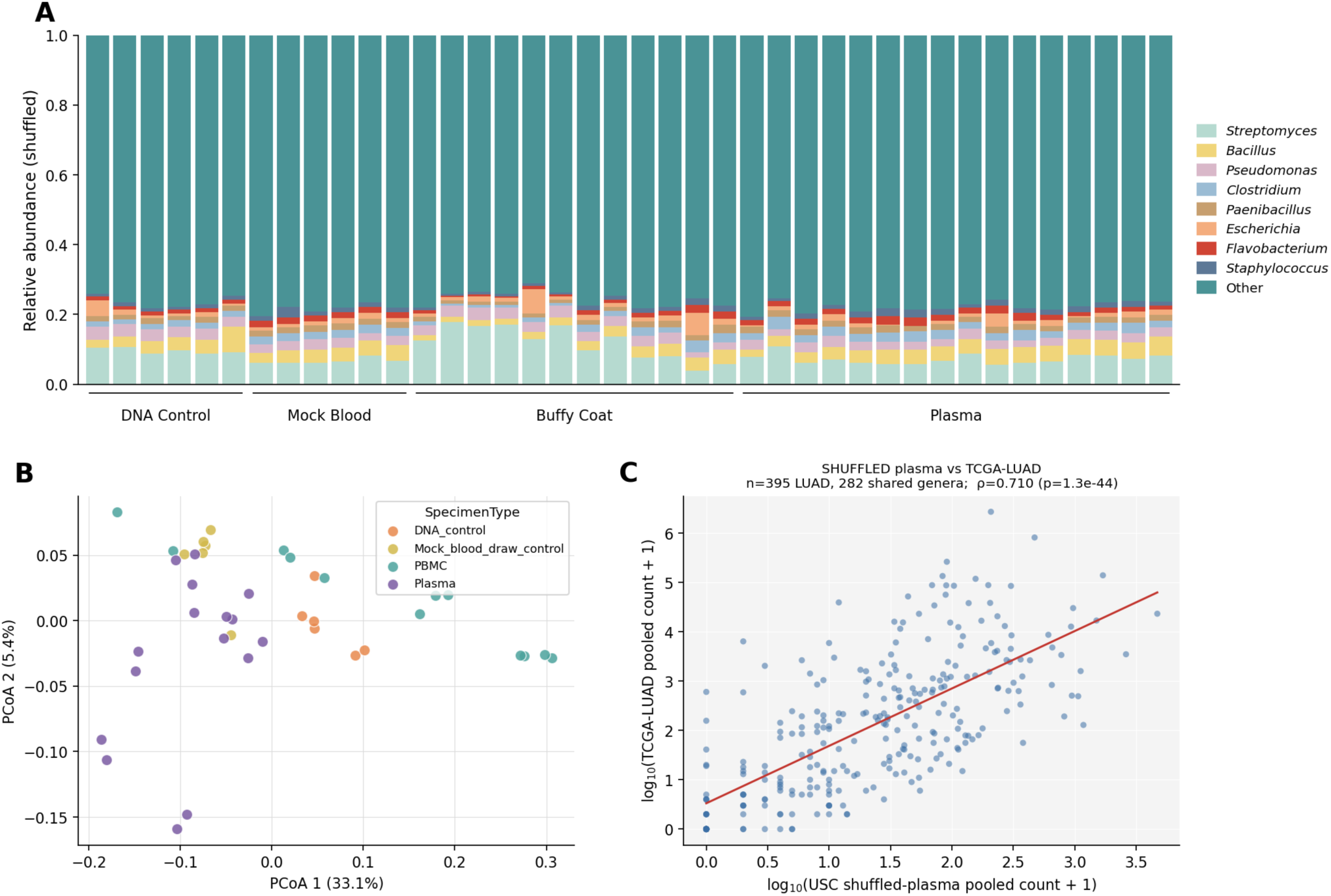
Structure, clustering, and cancer-cohort agreement persist after per-base shuffling (all panels on shuffled reads, which preserve GC content and length but randomize nucleotide order). (A) Stacked bars across the 40 shuffled libraries by specimen type, ordered by *Bacillus* share; the eight most abundant shuffled genera (*Streptomyces*, *Bacillus*, *Pseudomonas*, *Clostridium*, *Paenibacillus*, *Escherichia*, *Flavobacterium*, *Staphylococcus*) are colored, the rest pooled. (B) PCoA still separates by specimen type along PCoA 1 (33.1%), mirroring Figure 2. (C) Pooled shuffled-plasma versus TCGA-LUAD counts (n = 395) over 282 genera; ρ = 0.71, p = 1.3 × 10⁻⁴⁴; red line, OLS fit.

**Figure 4.**
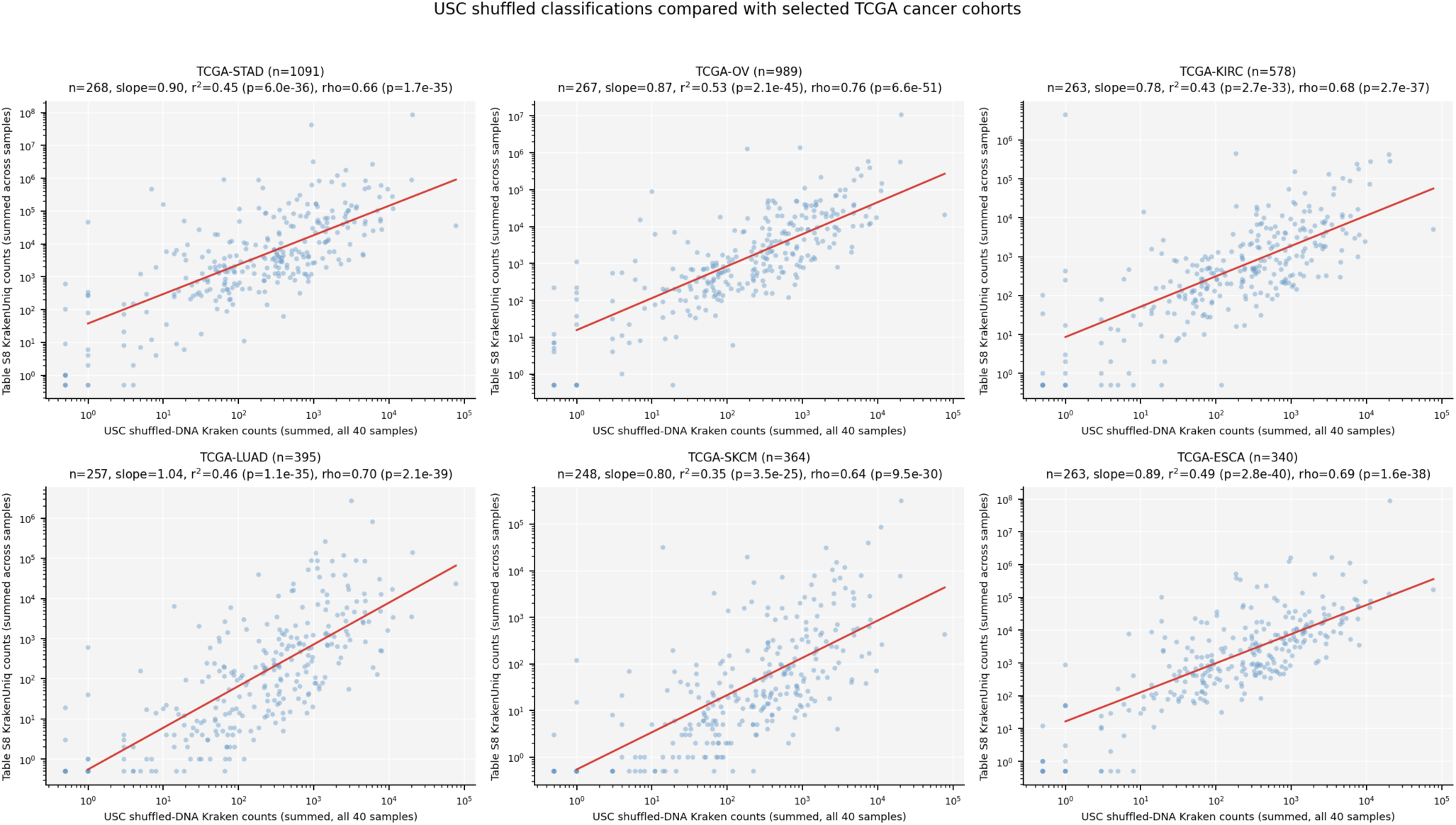
Classifications from previously published cancers (Sepich-Poore et al., Table S8) versus our 40 shuffled samples collapsed across libraries. Representative TCGA cohorts are shown; all per-cancer scatterplots are in Supplementary File 1.

That this correlation survives shuffling raises the question of how much of the shared signal reflects cancer biology at all. Comparing every sample group with the pooled TCGA-LUAD catalog (Table 1), all groups, including controls, retained strong correlation within a narrow band (unshuffled Pearson r = 0.74 to 0.84), with no group standing apart as biologically distinct. The agreement between our cmDNA profiles and the published catalog therefore does not depend on the biological provenance of the input DNA, pointing to a classification artifact rather than shared cancer biology.

**Table 1.** Correlation of each sample group’s pooled genus profile with the TCGA-LUAD pan-cancer microbiome catalog (Sepich-Poore et al., Table S8), before and after per-base shuffling.

| Study group | n samples | Unshuffled Pearson r | Shuffled Pearson r |
| --- | --- | --- | --- |
| Prostate cancer patient plasma | 6 | +0.800 | +0.673 |
| Prostate cancer patient PBMC | 6 | +0.812 | +0.744 |
| Healthy plasma | 10 | +0.786 | +0.695 |
| Healthy PBMC | 6 | +0.839 | +0.770 |
| Mock blood-draw control | 6 | +0.772 | +0.683 |
| SeraSeq reference-DNA control | 6 | +0.738 | +0.735 |
| All plasma | 16 | +0.792 | +0.696 |
| All PBMC (buffy coat) | 12 | +0.829 | +0.752 |
| All samples | 40 | +0.791 | +0.743 |

*PBMC, peripheral blood mononuclear cell. Every sample group correlates strongly with the published catalog both before and after shuffling, and the correlation does not track the biological provenance of the input DNA.*

### Per-genus read counts scale with reference-database k-mer representation

To find the cause, we asked whether database representation drives assignment, comparing per-genus Kraken2 counts with the unique k-mer seach genus contributes to the Kraken2 standard database (Figure 5). Pooled across all 40 samples, log-transformed unshuffled counts correlated strongly with database k-mer counts (Figure 5A; n = 5,464 genera, r² = 0.74, slope = 1.06, Spearman ρ = 0.91, p < 10⁻³⁰⁰), and the relationship tightened rather than collapsed after shuffling (Figure 5B; r² = 0.85, slope = 1.33, ρ = 0.94, p < 10⁻³⁰⁰). The same dependence appears in the independent catalog: pooling per-genus counts across all 7,827 TCGA samples in Sepich-Poore et al. Table S8 against the same k-mer counts gives a significant correlation over the 282 shared genera (Figure 5C; r² = 0.34, slope = 0.48, ρ = 0.59, p = 2 × 10⁻²⁷), despite no harmonization of pipeline or database. Reference-database structure therefore drives much of the classification in both datasets.

**Figure 5.**
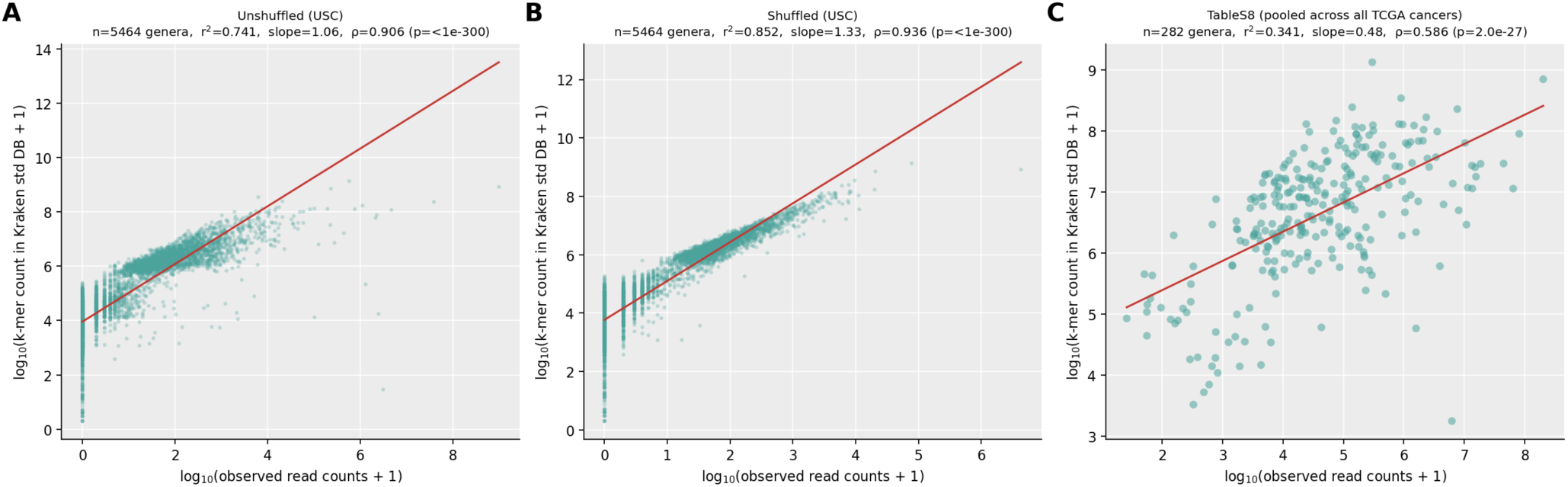
Per-genus read counts scale with the k-mers the Kraken2 standard database holds for each genus. Each point is a genus; x, log₁₀(observed reads + 1); y, log₁₀(database k-mers + 1); red lines, OLS fits. (A) Unshuffled reads, 40 samples (n = 5,464 genera; r² = 0.74, slope = 1.06, ρ = 0.91, p < 10⁻³⁰⁰). (B) Shuffled reads, same samples (r² = 0.85, slope = 1.33, ρ = 0.94, p < 10⁻³⁰⁰): the relationship tightens once biological sequence is destroyed, isolating database representation as the driver of assigned counts. (C) Pooled counts from Sepich-Poore et al. Table S8 (n = 7,827 samples) versus the same k-mer counts (n = 282 genera; r² = 0.34, slope = 0.48, ρ = 0.59, p = 2 × 10⁻²⁷).

### A shuffle-based negative control identifies 23 above-baseline taxa that collapse to a single defensible genus

We next used the shuffled assignments as a taxon-specific negative control, asking for each genus not merely whether a signal exists but whether it exceeds the shuffled baseline the same classifier produces. Regressing log-transformed unshuffled against shuffled per-genus counts across all genera (Figure 6), 23 taxa exceeded the shuffled-only model (Benjamini-Hochberg FDR ≤ 5%, two-sided); these are the taxa whose real-DNA signal is detectably larger than the artifact floor and for which a biological interpretation is potentially defensible.

**Figure 6.**
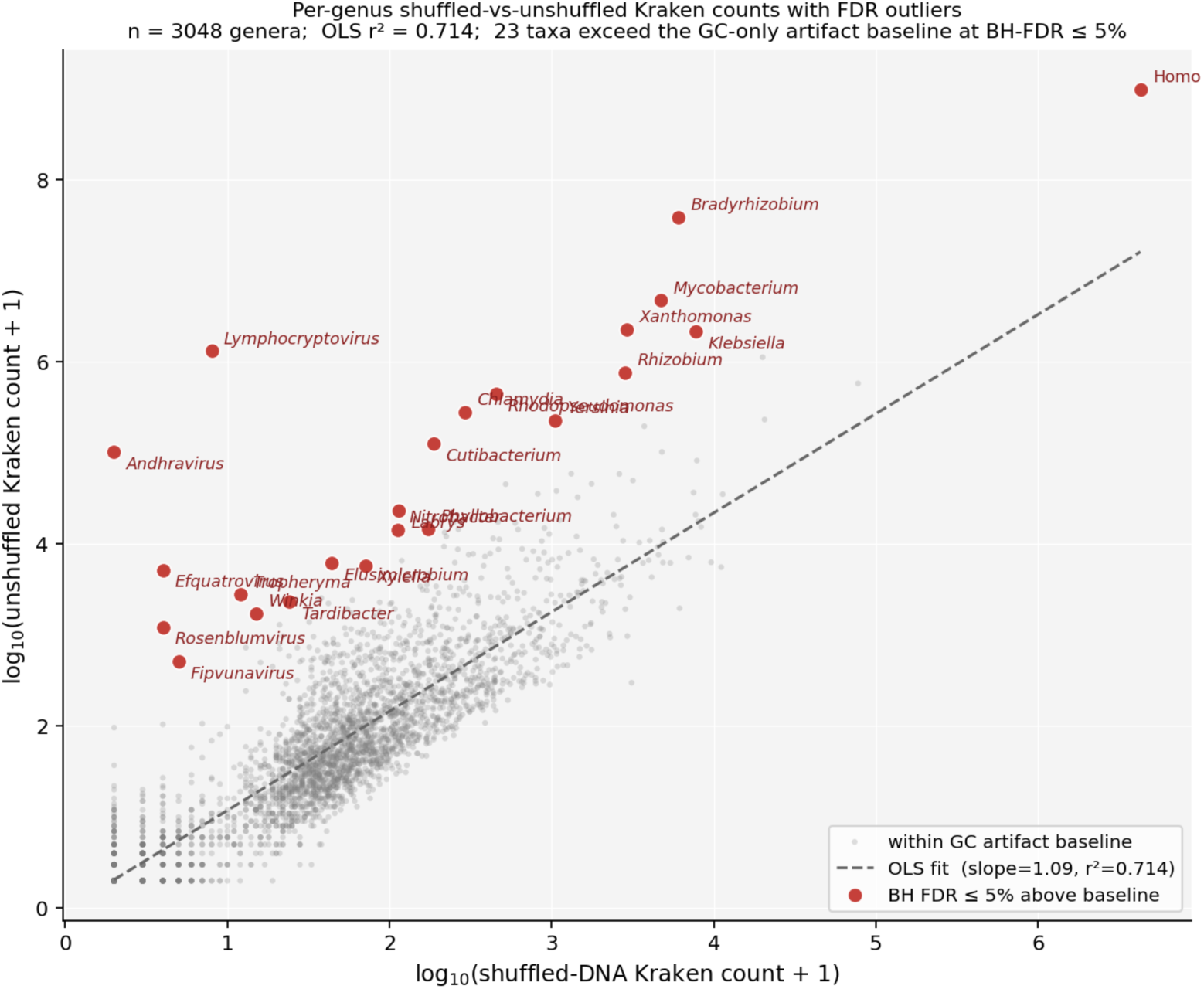
Per-genus log₁₀(shuffled Kraken2 count + 1) versus log₁₀(unshuffled count + 1) across all 3,048 Kraken2-reported genera. Grey, genera indistinguishable from the GC-only shuffled baseline; dashed line, OLS fit (slope ≈ 1.09, r² = 0.71); red points with italic labels, the 23 taxa exceeding the regression at Benjamini-Hochberg FDR ≤ 5% (Table 2).

**Table 2.** The 23 genera whose classified read counts exceed the GC-only shuffled baseline (Benjamini-Hochberg FDR ≤ 5%), grouped by likely origin.

| Category | n | Taxa (mean reads/sample) | n in MPA | Corroborated by MetaPhlAn |
| --- | --- | --- | --- | --- |
| Residual host DNA | 1 | *Homo* (24,562,081) | 0 | — |
| Known kit / reagent contaminants | 11 | *Bradyrhizobium* (968,818);<br>*Mycobacterium* (117,475);<br>*Xanthomonas* (56,517); *Rhizobium* (19,068);<br>*Rhodopseudomonas* (11,067);<br>*Cutibacterium* (3,140); *Labrys* (575);<br>*Phyllobacterium* (363); *Nitrobacter* (352);<br>*Elusimicrobium* (152); *Xylella* (145) | 3 | *Bradyrhizobium*,<br>*Cutibacterium*,<br>*Phyllobacterium* |
| Higher in non-patient (healthy or process) controls | 3 | *Lymphocryptovirus* (32,669);<br>*Andhravirus* (2,575);<br>*Efquatrovirus* (129) | 0 | — (MetaPhlAn database is bacteria-only) |
| Low abundance (<100 reads/sample) | 5 | *Tropheryma* (69);<br>*Tardibacter* (58);<br>*Winkia* (43);<br>*Rosenblumvirus* (30); *Fipvunavirus* (13) | 0 | — |
| Candidate signals surviving all filters | 3 | *Klebsiella* (53,526);<br>*Chlamydia* (6,945);<br>*Yersinia* (5,637) | 1 | *Klebsiella*<br>(*Chlamydia* and *Yersinia* not detected) |

MPA, MetaPhlAn. Counts are mean reads per sample across the 40 libraries. After removing host DNA, known contaminants, control-enriched taxa, and taxa below the <100-read power floor, and requiring orthogonal MetaPhlAn corroboration, only *Klebsiella* survives as a defensible circulating signal; this reflects survival of the filter, not a demonstrated cancer association.

We investigated the likely origin of each genus (Table 2). Eleven of twenty three are genera on published low-biomass contaminant lists [12,13] or established column- and reagent-borne background in plasma cmDNA: *Bradyrhizobium*, *Mycobacterium*, *Xanthomonas*, *Rhizobium*, *Rhodopseudomonas*, *Cutibacterium*, *Labrys*, *Phyllobacterium*, *Nitrobacter*, *Elusimicrobium*, and *Xylella*. *Bradyrhizobium*, the dominant buffy-coat taxon (Figure 2A), is consistent with contaminant DNA genuinely present but originating in sample handling. MetaPhlAn corroborates three of the eleven (*Bradyrhizobium*, *Cutibacterium*, *Phyllobacterium*), supporting real contaminant DNA rather than classification noise, consistent with their status as recognized skin- and reagent-derived contaminants [4,5,6]. Three viral genera, *Lymphocryptovirus* (Epstein-Barr virus, EBV), *Andhravirus*, and *Efquatrovirus*, distribute to SeraSeq DNA and mock blood-draw controls rather than patient plasma; because SeraSeq reference DNA carries EBV by design, EBV is disqualified as a patient signal, and none is callable by the bacteria-only MetaPhlAn database. Five further outliers (*Tropheryma*, *Tardibacter*, *Winkia*, *Rosenblumvirus*, *Fipvunavirus*) clear the threshold but at a low abundance averaging 13 to 69 reads per sample and all are unconfirmed by MetaPhlAn.

Three genera remain: *Klebsiella* (about 53,500 reads/sample), *Chlamydia* (about 6,900), and *Yersinia* (about 5,600) are the only high-abundance candidates. Of these, only *Klebsiella* is also called by MetaPhlAn; *Chlamydia* and *Yersinia* are absent from all 40 profiles. *Klebsiella* is therefore the single genus that passes all four validity criteria, (i) exceeding the GC-only shuffled baseline at FDR ≤ 5%, (ii) not appearing on a contaminant list, (iii) not being control-enriched, and (iv) being corroborated by an orthogonal marker-gene classifier. With only two patients and two volunteers, disease status is confounded with donor identity and no case-control test is feasible; the genus is detected by Kraken2 in all 40 samples, including the mock and reference-DNA controls (Table 3). MetaPhlAn was conservative, detecting no taxon in more than 12 of 40 samples (Table 3): it calls 13 genera, and Kraken2 calls all 12 NCBI-named one of these in nearly every sample (40 of 40 for 11, 39 of 40 for *Phyllobacterium*), whereas MetaPhlAn flags them in 1 to 12. The exception, GGB2722, is a MetaPhlAn species-level genome bin (SGB) with no NCBI name and no Kraken2 counterpart. The marker-gene method is thus strictly sparser: every genus it calls, Kraken2 also calls, in more samples, but Kraken2 calls thousands more that MetaPhlAn rejects. Whether MetaPhlAn conservative approach misses real low-abundance taxa warrants further work.

**Table 3.** The 13 genera detected by MetaPhlAn, with the number of samples (of 40) in which each was non-zero under each classifier.

| **Genus** | **MetaPhlAn (n non-zero)** | **Kraken2 (n non-zero, unshuffled)** |
| --- | --- | --- |
| Bradyrhizobium | 12 | 40 |
| Klebsiella | 12 | 40 |
| Cutibacterium | 10 | 40 |
| Acinetobacter | 4 | 40 |
| Brevundimonas | 4 | 40 |
| Methylobacterium | 4 | 40 |
| Sphingomonas | 4 | 40 |
| Bifidobacterium | 3 | 40 |
| GGB2722 | 3 | 0 |
| Phyllobacterium | 3 | 39 |
| Corynebacterium | 2 | 40 |
| Streptococcus | 2 | 40 |
| Pseudomonas | 1 | 40 |

*GGB2722 is a MetaPhlAn-internal species-level genome bin with no NCBI taxonomic name and therefore no Kraken2 counterpart. Every named genus MetaPhlAn calls is also called by Kraken2, in more samples.*

### Purely synthetic reads matched only to aggregate base composition and read length reproduce the misclassification

Every sample group, including pure SeraSeq DNA controls, mock blood draws, and the shuffled pool, correlated at ρ = 0.7 to 0.9 with TCGA-LUAD, consistent with classifier behavior rather than genuine biology. Because the shuffled sequences still use real reads as a base, we set a stricter bound by asking whether purely synthetic reads, generated de novo from a four-base random sampling process matched only to an aggregate GC target, still correlate with the published catalog. Although only 3,532 (0.0353%) of the 10⁷ synthetic pairs we generated received any Kraken2 assignment, the resulting Bracken profile correlated significantly with 27 of the 33 TCGA cohorts (FDR ≤ 5%, Spearman; Figure 7A); the six failing cohorts each had fewer than 20 samples (TCGA-ACC, -CHOL, -MESO, -PCPG, -THYM, -UCS). Among passing cohorts ρ ranged from 0.43 to 0.63, with TCGA-LUAD at 0.606 (p = 1.2 × 10⁻²⁹, BH q = 1.3 × 10⁻²⁸, slope = 1.64, R² = 0.34, n = 395, 282 genera; Figure 7B).

**Figure 7.**
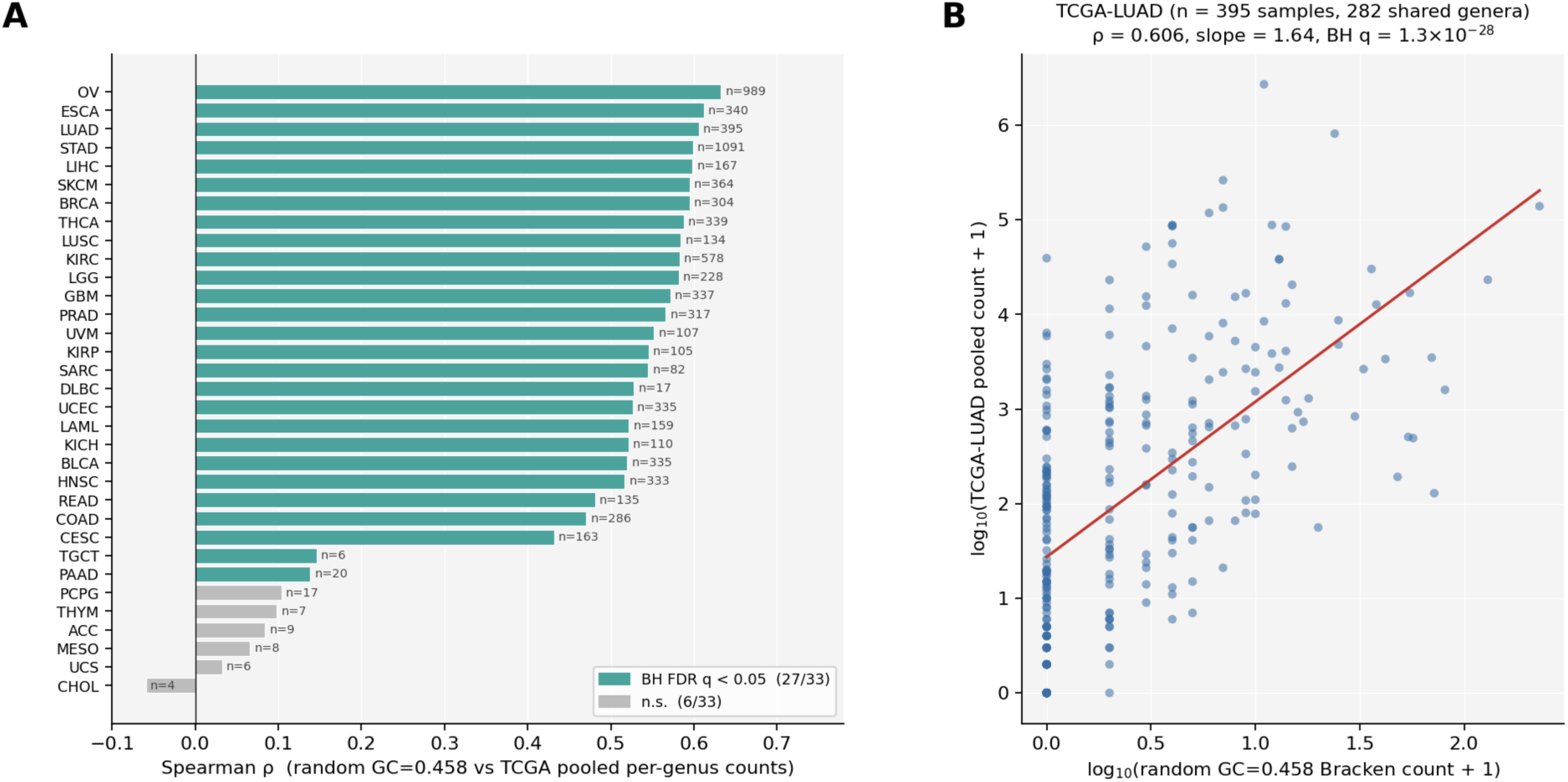
Synthetic reads at GC = 0.458, carrying no biological information, reproduce the cross-cohort agreement across most TCGA cancers. (A) Spearman ρ between the pooled Bracken profile of 10⁷ synthetic reads and each of the 33 TCGA cohorts (Sepich-Poore et al., Table S8) over 282 genera; bars colored by Benjamini-Hochberg FDR (teal, q < 0.05; grey, not significant); n, TCGA samples per cohort. (B) TCGA-LUAD example: each point is one of the 282 genera, log₁₀(synthetic count + 1) versus log₁₀(TCGA-LUAD count + 1); red line, OLS fit; ρ = 0.606, slope = 1.64, R² = 0.34, BH q = 1.3 × 10⁻²⁸.

This estimate is deliberately a floor. The shuffled plasma pool, preserving each read’s exact GC content across the library range (0.428 to 0.487, mean 0.483), gives ρ = 0.71 against TCGA-LUAD, and the unshuffled pool gives 0.81. The synthetic pool, with no per-sample GC variation, no strand asymmetry, and far fewer reads than any library (10⁷ versus 4 × 10⁶ to 2.7 × 10⁸ per sample), still yields a correlation of ρ = 0.61, versus 0.71 for the shuffled and 0.81 for the unshuffled pools, using only an aggregate base-composition target and a read-length distribution. Aggregate base composition and read length, processed through the Kraken2 database’s k-mer inventory, are therefore sufficient to manufacture a large fraction of the apparent agreement between cmDNA libraries and the published cancer-microbiome catalog independent of any other properties of the sequences.

### GC differences between samples distort statistical inference

Two plasma samples, one low-GC (45.8%) and one high-GC (48.4%), were each shuffled 20 times and reclassified (Figure 8, left and middle). Comparing the two directly (Figure 8, right), the lower-GC sample had higher classification rates across all taxa, so GC differences alone drive systematic classification differences even among shuffles of one base sample. Within shuffles from a sample, the null hypothesis of no difference should always be true. Two-sample tests within the low-GC (Figure 9, left) or high-GC (Figure 9, middle) shuffles gave uniform p-values, with about 5% below 0.05. By contrast, comparing taxonomic classifications from high-versus low-GC shuffles gave p-values far below the null, with nearly half of taxa differing despite all being based on shuffled sequences (Figure 9, right). These results demonstrate that even modest differences in mean GC content can generate many spurious differentially abundant taxa on non-biological shuffled sequences.

**Figure 8.**
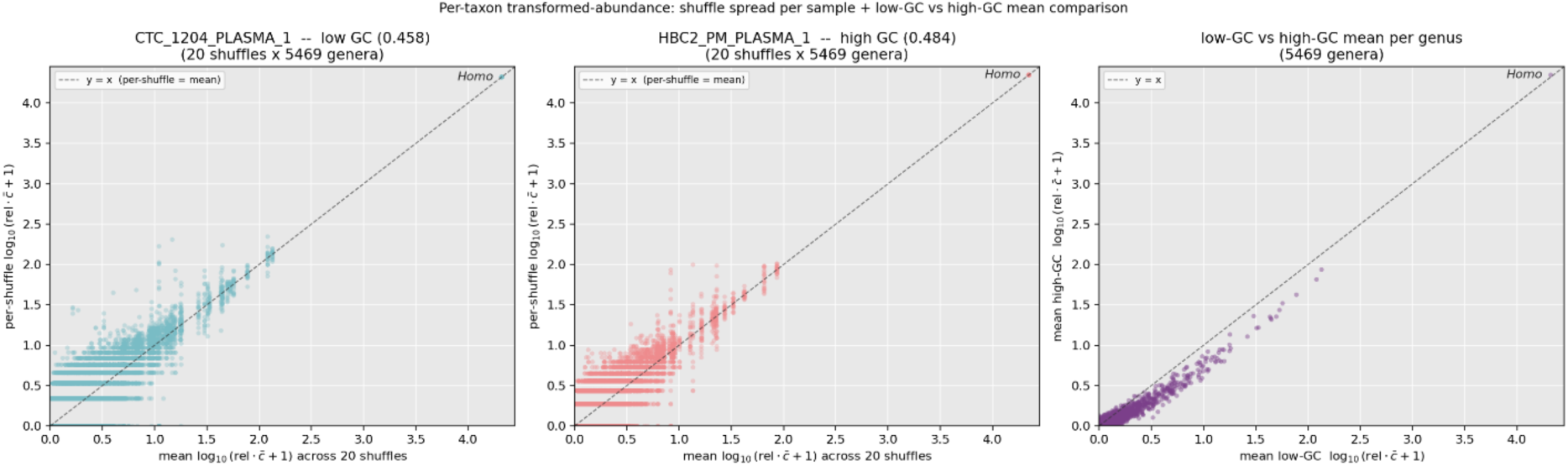
Per-taxon relative-abundance spread across 20 per-base shuffles of two plasma samples spanning the GC range (left, low-GC 45.8%; middle, high-GC 48.4%; right, high-versus low-GC). Each point is one shuffle of one genus; x, per-shuffle log₁₀ relative abundance; y, genus mean across the 20 shuffles. Horizontal spread at fixed y is the artifact noise floor: precisely counted taxa track the y = x diagonal, while rare taxa fan out as their per-shuffle counts swing across orders of magnitude.

**Figure 9.**
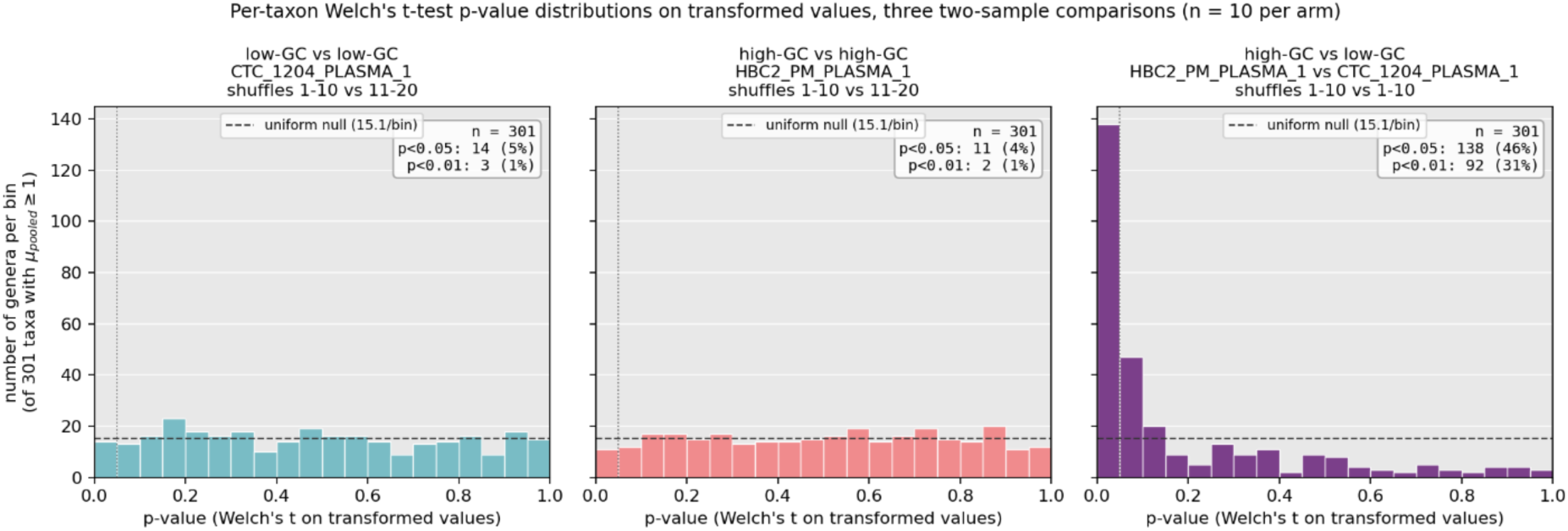
Per-taxon Welch’s t-test p-value distributions for three comparisons among shuffle replicates of the two plasma samples (low-GC 45.8%, high-GC 48.4%; 20 shuffles each), over every genus with non-trivial counts (mean pooled ≥ 1 read; n = 301) on the log-transformed abundance scale; 10 shuffles per arm to equalize power; dashed line, uniform null. The two within-sample panels track the null (4.7% and 3.7% of taxa at p < 0.05; medians 0.473 and 0.516). The cross-GC panel is sharply non-uniform (45.8% at p < 0.05, 30.6% at p < 0.01; median 0.061), with a spike near zero. With sample size held constant, this reflects GC-driven artifact rather than differential power.

## Discussion

Our findings reveal several dangers in analyzing low-biomass clinical samples with out-of-the-box taxonomic pipelines. In our initial analysis of our 40 samples, buffy coats and plasma differed markedly, with buffy coats dominated by *Bradyrhizobium* and plasma and controls largely human even after host-read removal, producing clear PCoA clustering that could be read as biology. Moreover, pooled Kraken2 counts agreed strongly with a previously published dataset suggesting robust cross-dataset reproducibility. Yet PCoA on shuffled sequences still separates by specimen type, and shuffled classifications still correlate with the published cancer data despite no pipeline harmonization. One driver is how each taxon is represented in the reference database, evident when regressing observed counts against Kraken’s k-mer database representation. Together these analyses indicate that much of the structure in this dataset can be explained by pipeline properties rather than biology.

Both informatic and laboratory controls are therefore essential when biomass is low. Here, plasma had matched DNA and mock-draw controls but PBMCs did not, leaving *Bradyrhizobium* without a data-driven basis for exclusion; specimen-type-specific negative controls following identical wet-lab workflows would enable objective contaminant identification. Classification also needs scrutiny: through shuffled-sequence regressions, k-mer matching, and fixed-GC synthetic reads, we show Kraken2 is prone to systematic misclassification driven by the reference database. MetaPhlAn’s marker-gene approach is more conservative, giving fewer but more trustworthy calls at the risk of missing sparse signal [16]; where Kraken2’s breadth is wanted, a second method should corroborate it.

We note that Kraken2 does not store k-mers directly. For each 35-mer in a reference genome it computes a 31-bp minimizer under a scrambled canonical ordering, and the database is a compact hash table keyed on those minimizers, each mapping to the lowest common ancestor of all taxa whose k-mers share it. Many distinct k-mers therefore can collapse to one stored entry, and the per-genus counts used here are counts of database entries attributed to a genus — minimizers, not k-mers. We retain the term k-mer throughout because it names the quantity the classifier is nominally estimating and because the compression is monotone in genome content, but two consequences of the minimizer scheme bear on the interpretation of Figure 5. First, the compression ratio is not uniform across taxa: it depends on repeat structure and base composition, so a genus’s stored representation is a noisy proxy for its true k-mer content.

Second, and more directly relevant, minimizer selection is itself a function of local base composition, so the compositional channel that per-base shuffling exposes operates through minimizer selection rather than around it. The relationship in Figure 5 is thus best read as a dependence on how densely the database is populated for each genus, whatever the underlying algorithmic cause.

We note that only 0.0353% of the purely synthetic pairs received taxonomic classification. However, in a standard bioinformatics pipeline, ordination, cross-cohort correlation, and per-taxon differential abundance are computed on relative abundances which are invariant to overall classified depth. The 3,532 classified read-pairs in the synthetic experiment, carrying no information beyond an aggregate GC target, were sufficient to recover a Spearman ρ of 0.61 against TCGA-LUAD across 282 shared genera, most of the ρ = 0.81 obtained from the real libraries. So even a very low classification rate yields a pattern of taxa that reflects the signal caused by the distribution of k-mers in the reference database rather than biological signal. In a Kraken database, the number of bits that are reserved for avoidance of hash collision can vary depending on the number of taxa in the database and this is one of the factors that can influence the fraction of shuffled reads that will be classified. But even where fewer or no shuffled sequences are classified, the prior distribution of k-mers in the database should be explicitly considered when interpreting low-abundance signal

A partial remedy for the issues raised here is available within Kraken2 itself. The classifier can be run to filter by confidence score — the fraction of a read’s minimizers mapping to the assigned taxon’s clade — and raising the threshold above the default of zero discards reads whose support is spread thinly across the taxonomy, which is the signature of the compositional matches described here. But thresholding is not a complete solution as there is no principled procedure for choosing the value: confidence trades false positives against false negatives, and calibrating that trade requires knowing the sensitivity cost, which in turn requires a positive control of defined composition that low-biomass studies rarely include. A threshold high enough to suppress compositional artifact in one library may discard authentic low-abundance signal in another, and nothing in the output distinguishes the two cases. We did not utilize a confidence score cutoff in this manuscript. Our work suggests as an alternative with shuffled control sequences offering a way to filter empirically rather than by an arbitrarily chosen threshold.

Because shuffled reads contain no biological sequence by construction, every assignment they receive is a false positive and the distribution of shuffled reads can therefore determine where the artifact floor is. This is the basis of the regression in Figure 6, which retained 23 genera above that floor from several thousand reported by Kraken2. While shuffling can distinguish real sequence from GC- and database-driven artifact, it will not identify contaminating DNA; a taxon above the shuffled null is established as sequence that exists in the dataset, not as biologically meaningful, and discriminating it from a contaminant still needs an external prior such as informed comparison to a database of known kit contaminants.

Most circulating microbiome studies remove human reads before taxonomic classification, assuming that the remaining sequences represent microbial DNA rather than host DNA. Our findings indicate that, although necessary, this safeguard alone is insufficient. Per-base shuffling preserves each read’s GC content while destroying every meaningful k-mer, yet shuffled libraries still yield tens of thousands of genus assignments per sample, still cluster by specimen type, and still correlate at ρ ≈ 0.7 to 0.9 with existing cancer datasets, a correlation devoid of biological information by construction. The mechanism is direct: after host removal, what remains is a GC distribution, and any database whose k-mer inventory is itself structured by genome GC content (as the Kraken2 standard database is; Figure 5) assigns those reads to taxa in the matching compositional band. Host-read removal drops sequences matching the human reference but not sequences that merely share base composition with microbial k-mers, and it is that compositional channel the shuffle exposes. Host depletion should therefore be paired with a GC-only negative control, most simply per-base shuffling of the same reads, and candidate taxa required to exceed it before interpretation. Purely synthetic reads matched to aggregate base composition and read length, which reproduced much of the cross-cohort agreement while containing no organisms and no preserved sequence, set a hard lower bound on how much of that agreement could be biological.

The GC-content experiment also has important implications for differential abundance testing. Within-sample comparisons behaved correctly, with uniform p-values and about 5% significant, whereas the cross-GC comparison was sharply anti-conservative, with nearly half of genera significant despite no real difference. Because GC content varies across the genome and is perturbed by tumor copy-number alterations and coverage bias, cases and controls differing even modestly in mean GC can generate many spurious differentially abundant taxa. GC matching, or explicit GC adjustment in the model, is therefore a prerequisite for valid differential-abundance testing in low-biomass cmDNA studies, and the shuffle-derived floor gives a principled baseline for the effect sizes such studies must be powered to detect.

Synthesizing these results, we propose a four-criterion validity test to apply per taxon: a Kraken2 cancer signal is credible only if it (i) lies significantly above the same classifier’s output on per-base-shuffled reads from the same libraries, (ii) is not on an established kit or reagent contaminant list, (iii) is not enriched in non-patient controls, and (iv) is reported by at least one orthogonal classifier such as MetaPhlAn. Applied to our catalog of several thousand genera, this reduces the candidates to a single genus, *Klebsiella*, showing how much apparent cmDNA biology is a property of the classifier rather than the sample. In addition, comparison of samples with different GC content can produce apparent differences in taxonomic composition that are independent of sample biology, so care must be taken to avoid over interpreting inference in such cases.

This study is a single-center pilot with few patients and volunteers, so it characterizes contamination and artifact with strong internal control but cannot estimate population-level prevalence or prove the absence of an authentic signal. Analyses are genus-level; one reference cohort (TCGA-LUAD) is the running example, though the pattern holds across most of the 33 TCGA cohorts; results are specific to the Kraken2 standard database build tested; no positive control, defined-composition mock community, or spike-in standard was run through the pipeline, so the sensitivity (false-negative rate) of the validity filter and any absolute quantification remain uncharacterized; and because the MetaPhlAn database is bacteria-only, the orthogonal-corroboration criterion cannot be satisfied by any virus, disqualifying viral candidates for a structural rather than biological reason. Even so, a single defensible genus emerging from thousands, and the reproduction of the published signal from pure noise, argue that rigorous negative controls must precede any biological claim.

## Conclusions

In this controlled pilot, much of the apparent cmDNA signal in blood, including its agreement with a published pan-cancer catalog, is explained by base composition and reference-database architecture rather than authentic biology, and short-read k-mer pipelines cannot separate the two on their own. Low-biomass cmDNA studies should adopt specimen-matched negative controls, a conservative second classifier, per-base shuffling (with GC-matched synthetic reads as a stricter floor), and GC-aware analysis and design, and candidate taxa should clear these controls before being advanced as cancer biomarkers.

## Acknowledgements

We would like to thank and acknowledge the participants who generously donated their biospecimens for study. This work was supported in part by the Engineering Research Centers Program of the National Science Foundation under NSF Cooperative Agreement No. EEC-2133504 and by the Department of Bioinformatics and Genomics at UNC Charlotte. Work conducted by Amir Goldkorn and Daniel Bsteh has been supported in part by funds from the Rivals United for a Kure Grant (Kure It Foundation) and P30CA014089 (NCI/NIH). Wet lab contributions to this project were executed by Daniel Bsteh, while computational analysis was executed by Daisy Fry Brumit and Anthony Fodor. Code testing and review was completed by Shan Sun, Daisy Fry Brumit, and Anthony Fodor, with contributions by Ivory Blake. Writing was executed by Daisy Fry Brumit, Daniel Bsteh, Anthony Fodor, and Tanya Alderete, with significant edits and contributions by Amir Goldkorn, Jesse Goodrich, and Michael Liss.

## COI Disclosure

Authors report no conflicts of interest with this work.

*All figures (Graphical Abstract, Figures 1-9) and tables (Tables 1-3) with legends are provided in the accompanying file, “Circulation Microbial DNA Figures and Tables.”*

